# Insights into the effect of the J-domain on the substrate binding domain (SBD) of the 70 kDa heat shock protein, Hsp70, from a chimeric human J-SBD polypeptide

**DOI:** 10.1101/251900

**Authors:** Ana O. Tiroli-Cepeda, Thiago V. Seraphim, Júlio C. Borges, Carlos H. I. Ramos

**Affiliations:** Universidade Estadual de Campinas, Campinas SP, 13083-970, Brazil; São Carlos Institute of Chemistry, University of São Paulo, São Carlos, SP, 13560-970, Brasil

**Keywords:** chaperone, Hsp70, Hsp40, J-domain, Hsp70-SBD, chimera, protein-protein interaction

## Abstract

DnaJ/Hsp40 chaperones deliver unfolded proteins and stimulate the ATPase activity of DnaK/Hsp70 via their J-domain, a crucial event for the function that this system has in assisting protein folding. The interaction between Hsp40 and Hsp70 is transient and thus difficult to study, since mixing the binding partners can lead to quick dissociation due to their low affinity, creating a challenge for detailed analysis. As a consequence, knowledge of many important aspects of the mechanism of interaction is still lacking, for instance, the effect that J-domain binding has on Hsp70. In this study, we investigated whether it would be possible to gain understanding of this interaction by engineering a chimeric polypeptide where the J-domain of Hsp40 was covalently attached to the substrate binding domain (SBD) of Hsp70 by a flexible linker. The rationale for this is that an increase in the proximity between the interacting partners in this engineered chimera will promote the natural interaction and facilitate the characterization of the protein– protein interaction, which is a requirement to gain further understanding of many biological processes. The resulting chimera, termed J-SBD, was properly folded and had properties not present in the SBD alone. J-SBD behaved primarily as a monomer in all conditions tested and exhibited chaperone activity, as shown by aggregation protection and substrate binding assays, which revealed decreased binding to bis-ANS, a probe for hydrophobic patches. Collectively, our results suggest that Hsp40 binding to Hsp70 via the J-domain shifts the Hsp70 equilibrium towards the monomer state to expose hydrophobic sites prone to substrate accommodation.

**Abbreviations:** Bis-ANS (4,4’-Dianilino-1,1’-Binaphthyl-5,5’-Disulfonic Acid; CD, circular dichroism; Hsp, heat shock protein; J-SBD, chimeric polypeptide in which the J-domain of Hsp40 (at the N-terminus) is covalently attached to the substrate binding domain of Hsp70 (at the C-terminus) by a flexible linker; SBD: substrate binding domain of Hsp70.

## 1. Introduction

Proteins are involved in a large array of cell functions, and to be able to perform these functions, these macromolecules must be properly folded. As one of several systems that aid folding inside the cell, the Hsp70 chaperone system is of great significance, since it plays an essential role in protein homeostasis (Mayer et al, 2001; Hartl & Hayer-Hartl, 2002; Kampinga & Craig, 2010; da Silva & Borges, 2011; Tiroli-Cepeda & Ramos, 2011; Pryia et al, 2013; Alderson et al, 2016). As a matter of fact, the Hsp70 system has been reported to be involved in preventing aggregation, assisting folding, participating in transport across membranes, and many other functions.

Proteins from the Hsp70 family contain two domains. The Nucleotide Binding Domain (NBD) is located at the N-terminus, is approximately 45 kDa, and has ATPase activity (Chappell et al, 1987; Flaherty et al, 1990). The substrate binding domain (SBD; Fig. 1A) is located at the C-terminus, is approximately 25 kDa and is involved in binding the polypeptide substrate (Wang et al, 1993; Morghauser et al, 1995). The conformation of the SBD is under the control of the NBD because when there is ATP in the NBD, the SBD is mainly found in an open conformation and has a low affinity for substrate. On the other hand, when the ATP is hydrolyzed, the SBD mainly assumes a closed conformation and binds the substrate with higher affinity, which is a consequence of the structural changes promoted.

**Fig. 1:**
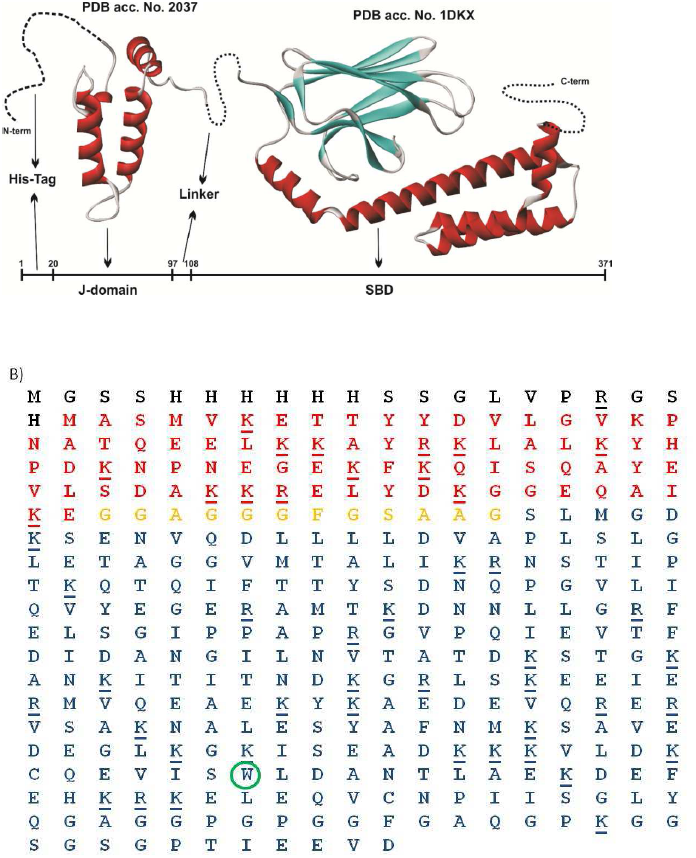
SBD and chimeric J-SBD. A) Schematic representation of structural domains used in the construction of the chimera. PDB 2O37: J-domain of Sis1 from *Saccharomyces cerevisiae*. PDB 1DKX: substrate binding domain of DnaK from *Escherichia coli*. B) Sequence of the chimeric J-SBD, which contains the J-domain of human Hsp40-1A (red), connected to the SBD of human Hsp70-1A (blue) by a flexible linker (yellow). The J-SBD protein was produced with a histidine-tag (black). Lysine and arginine residues are underlined and the single tryptophan residue is highlighted.

The chaperone activity of Hsp70 is regulated by its co-chaperones, such as those from the DnaJ/Hsp40 family (Szabo et al, 1994; Minami et al, 1996; Summers et al, 2009; Cyr & Ramos, 2015). The ATPase activity of Hsp70 is stimulated by Hsp40, and at the same time this co-chaperone delivers the substrate, or client protein, to Hsp70 (Mayer et al, 2001; Summers et al, 2009). Therefore, Hsp40 is critical for efficient activity of Hsp70, acting as a holder (i.e., recognizing and binding non-native polypeptides to prevent aggregation) (Lu and Cyr, 1998; Summers et al, 2009; Cyr & Ramos, 2015). The presence of the J-domain, which is approximately 70 amino acid residues long, characterizes Hsp40 co-chaperones. The J-domain is highly conserved and, in the cases of types I and II Hsp40s, is at the N-terminus (for reviews see Kampinga & Craig, 2010; Cyr and Ramos, 2015). The J-domain triggers Hsp70 ATP hydrolysis and locks the chaperone onto the client protein, thus promoting its folding (Fan et al, 2003; Walsh et al., 2004). In agreement with this function, although isolated co-expression of either Hsp40 or Hsp70 in O23 hamster cells is sufficient to protect luciferase from aggregation, only the combined co-expression of both Hsp40 and Hsp70 promotes the refolding of luciferase (Annemieke et al., 1997).

Therefore, to further understand the mechanism by which Hsp70 aids folding, it is important to understand the effects of Hsp70 interaction with Hsp40. However, this task is not easy since the interaction is transient. Hsp40 binds Hsp70 and dissociates as fast as the ATP is hydrolyzed (Misselwitz et al., 1999), and this is dependent on the N-terminal domain of Hsp70 (Gässler et al 1998). Moreover, the mechanistic roles of J-proteins in regulating Hsp70 function and how the J-protein impacts Hsp70-substrate complex formation are not yet very well understood. To find possible clues that might help increase our knowledge about the interaction of Hsp70 with Hsp40, we engineered a chimeric polypeptide using molecular biology tools, in which the SBD of Hsp70 is covalently attached to the J-domain by a flexible linker. The rationale of creating this chimera is that an increase in the proximity between the interacting partners will promote their natural interaction and facilitate characterization of this protein–protein interaction, which is a requirement to gain understanding of biological processes. The resulting chimera, termed J-SBD, was properly folded and was characterized by several biophysical tools. The chimeric J-SBD showed properties not present in the SBD alone: it behaved as a monomer in all conditions tested, exhibited chaperone activity as showed by luciferase aggregation protection assay and was capable of binding substrates, as shown by decreasing binding to bis-ANS, a probe for hydrophobic patches. Collectively, our results suggest that Hsp40 binding to Hsp70 via the J-domain shifts the Hsp70 equilibrium towards the monomer state to expose hydrophobic sites prone to accommodate substrates.

## 2. Materials and Methods

### 2.1. Cloning, expression and purification

The DNA fragment coding for the J-domain of human Hsp40 (DjA1/Hdj2/dj2/HSDJ/Rdj1 from subfamily A) was amplified by PCR from the pET28aDjA1 plasmid (Borges et al., 2005) using the forward primer 5’ CCGGCAG**GCTAGC**ATGGTGAAAGAAACAAC 3’ (primer 1), which introduced a *Nhe*I restriction site, and the reverse primer 5‵ATCAAA**GGATCC**CGCGGCGGAG 3‵ (primer 2), which introduced a *Bam*HI restriction site. The DNA fragment coding for the SBD of human Hsp70-1A I was amplified by PCR from the pET28aHsp70-1A plasmid (Borges et al, 2006) using the forward primer 5‵GCAGGCG**GGATCC**CTATGG 3’ (primer 3), which introduced a *Bam*HI restriction site, and the reverse primer 5‵ATCAAA**GGATCC**CGCGGCGGAG 3‵ (primer 4), which also introduced a *Bam*HI restriction site. Then, the two PCR products were digested with *Bam*HI and ligated to generate chimeric cDNA containing the sequence of the J-domain of Hsp40 fused to the SDB domain of Hsp70. The chimeric cDNA was then digested with *Nhe*I and cloned into a pET28a expression vector. The correct cloning was confirmed by DNA sequencing using an ABI 377 Prism system (PerkinElmer Life Sciences). These procedures created the pET28aJ-SBD vector, which was transformed in *E. coli* strain BL21(DE3) for protein expression by adding 0.4 mM isopropyl thio-b-D-galactoside at *A*_600_ ~0.8 AU. The induced cells were grown for 3 h at 37 °C, and harvested by centrifugation for 10 min at 2600 x *g*. The bacterial pellet was resuspended in 25 mM Tris-HCl, pH 8.0, and 150 mM NaCl (15 mL L^-1^ medium), disrupted by sonication in an ice bath, and centrifuged for 15 min at 12000 x *g*. The supernatant was subjected to metal affinity chromatography in a HiTrap Chelating column (Amersham Biosciences), using an AKTA FPLC (Amersham Biosciences). The J-SBD chimeric polypeptide was eluted in 25 mM Tris-HCl (pH 8.0) buffer containing 150 mM NaCl and 500 mM imidazole and loaded onto a HiLoad Superdex 200 pg molecular exclusion column using an AKTA FPLC. The degree of purification was estimated by SDS-PAGE, and protein concentration was determined spectrophotometrically, using the calculated extinction coefficient for denatured proteins (Edelhoch, 1967; Ramos, 2004). All buffers used were of chemical grade and were filtered before their use to avoid scattering from small particles. The pPROEXSBD vector, containing the SBD domain of human Hsp70-1A, was a gift from Dr. Jason Young (McGill University, Canada), and the procedures of expression and purification for the JSBD chimera were the same as those described above. Unless otherwise stated, buffer conditions for protein study were 25 mM Tris-HCl (pH 8.0) and 150 mM NaCl.

### 2.2. Spectroscopy

Circular dichroism (CD) spectra were recorded in a JASCO model J-810 CD spectropolarimeter equipped with a thermoelectric sample temperature controller (Peltier system) using standard conditions (Correa & Ramos, 2009). Data were collected from 260 nm to 200 nm and were averaged at least ten times. All measurements were made in cuvettes with a 1 mm pathlength and with a protein concentration of 10 μM.

Emission fluorescence spectra were recorded on an Aminco Bowman^®^ Series 2 (SLM-AMINCO) fluorimeter using quartz cells of 10 x 10 mm optical path length. Protein concentrations were 5 μM. Emission fluorescence spectra of tryptophan were obtained with excitation at 295 nm (bandpass of 4 nm) and with emission from 304 to 400 nm (bandpass of 4 nm). Emission fluorescence data were analyzed either by their maxima wavelength or by their center of spectral mass (<λ>), as described by the equation below:

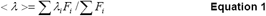

where λ*_i_* is each wavelength and *F_i_* is the fluorescence intensity at λ*_i_* (Silva et al., 1986). All data were analyzed with Origin software (Microcal).

Bis-ANS (4,4’-Dianilino-1,1’-Binaphthyl-5,5’-Disulfonic Acid) was incubated with 2 μM protein, at a concentration of 20 μM for 10 min at 25 °C, and emission fluorescence spectra were measured with excitation at 365 nm (bandpass of 4 nm) and emission from 400 to 600 nm (bandpass of 4 nm). Experiments were performed in the absence or in the presence of peptide (NRLLLTG) and absorbance at 495 nm was reported. Each curve was an average of at least three independent experiments.

### 2.3. SEC-MALS

The oligomeric states of SBD and J-SBD chimeric protein were investigated by size exclusion chromatography coupled with multi-angle laser light scattering (SECMALS). Experiments were performed using the Superdex 200 10/300 GL column (GE Life Sciences) and 100 μL of samples (2 and 4 mg mL^-1^ for J-SBD and 2, 4 and 9 mg mL^-1^ for SBD) were injected onto the column. Light scattering measurements of the proteins eluted from the column were performed with a miniDAWN TREOS detector (Wyatt Technologies). Data analysis and molecular mass values were obtained using ASTRA software (Wyatt Technologies).

### 2.4. Analytical ultracentrifugation

Sedimentation velocity (SV) analytical centrifugation experiments were carried out in a Beckman Optima XL-A analytical ultracentrifuge using AN-60TI rotor using standard conditions (Borges & Ramos, 2011). The J-SBD protein was prepared in 50 mM Tris-HCl (pH 7.5) buffer containing 100 mM NaCl at concentrations ranging from 0.15 to 1.0 mg mL^-1^. The SV experiments were carried out at 20 °C, 35,000 rpm and data acquisition was performed at 229 nm and 236 nm. The software SedFit (Version 12.1) was used for data treatment. This software fits the absorbance versus cell radius data and models the Lamm type equation to discriminate spreading of the sedimentation boundary from the diffusion function, supplying the continuous sedimentation coefficients c(S) distributions (Schuck, 2000),. As a regularization parameter, weight average value of frictional ratio (*f*l/*f*l_0_) was allowed to float freely. The software Sednterp was used to estimate viscosity (η = 0.010267 poise) and density (ρ = 1.00379 g mL^-1^) for buffer at 20 °C and to estimate the Vbar (0.7307 mL g^-1^) for the J-SBD protein. These parameters helped to correct the experimental s-value obtained from the peak of the c(S) distribution at each protein concentration and s_20,w_ (the sedimentation coefficient at standard conditions (water and 20 °C)) and the standard sedimentation coefficient at infinite dilution (0 mg mL^-1^, s^0^_20,w_), by linear regression, from the curve of s_20,w_ versus protein concentration. The value of s^0^_20,w_ is an intrinsic property of the protein and contains information about the molecular mass (MM) and asymmetry of the particle.

### 2.5. Aggregation protection

Luciferase (Sigma) 1 μM was heated to 42 °C in a 1 cm quartz cuvette in the absence or presence of J-SBD (1 to 10 μM, see figure caption for details), in 25 mM Tris-HCl (pH 8.0) buffer containing 150 mM NaCl. Scattering (turbidity) was followed at 320 nm for 600 s in a spectrophotometer (Jasco UV/VIS 530). Each curve was an average of at least three independent experiments.

### 2.5. Protease accessibility assay

Each protein was incubated at 20 μM concentration with trypsin in a 1:200 ratio (protease:protein) at 20 °C and was sampled at 5, 10, 15, 20, 30, 40, 50, 60, 90, 120 and 180 min. The reactions were stopped with 1% TFA. Digestions were analyzed using SDS-PAGE. Samples at 20, 60 and 180 min were submitted to an affinity assay on Ni-NTA sepharose, with an incubation time of 15 min, followed by three washes with 25 mM Tris-HCl (pH 8.0) containing 150 mM NaCl, and three elutions with 25 mM Tris-HCl (pH 8.0), 200 mM imidazole, and 150 mM NaCl.

## 3. Results and Discussion

### 3.1. The chimeric J-SBD was produced folded

Hsp70 and Hsp40 are major cytosolic molecular chaperones that aid protein refolding and prevent unspecific protein interactions that could otherwise lead to aggregation by Hsp70-dependent ATP hydrolysis (for reviews see Kampinga & Craig, 2010; Tiroli-Cepeda & Ramos, 2011; Priya et al., 2013; Cyr & Ramos, 2015; Alderson et al, 2016). When overexpressed, these chaperones increase cell resistance to stress and suppress degenerative phenotypes. The interaction cycle starts with Hsp40 delivering unfolded proteins to Hsp70 and, via its J-domain, stimulating ATPase activity, an event crucial to aiding protein folding. The interaction between Hsp40 and Hsp70 is transient and thus difficult to study because mixing the binding partners can lead to quick dissociation due to their low affinity, challenging detailed analysis. The J-domain interacts mainly with the NBD and the flexible linker of Hsp70, but it has been shown that the SBD interacts with both Hsp40 and TPR-proteins in an independent manner (Demand et al, 1998). However, detailed aspects of the mechanism of interaction are still lacking, for instance, the effect that the binding of the J-domain has on Hsp70 substrate binding domain (SBD).

We investigated whether it would be possible to increase current knowledge about the interaction between Hsp70 and Hsp40 by engineering a chimeric polypeptide, in which the J-domain is covalently attached to the SBD by a flexible linker (Fig. 1). The rationale for this experiment is that an increase in proximity between the interacting partners in the chimera will promote the natural interaction and facilitate the characterization of protein–protein interactions, which is a requirement to gain an understanding of biological processes. Fig. 1B shows the sequence of the chimeric protein J-SBD, which contains the J domain of human Hsp40 fused to the SBD of human Hsp70, produced with a His-tag at the N-terminus. The resultant chimera, termed J-SBD, was produced folded and had its conformation characterized using several biophysical tools (Batista et al, 2015), as described below.

Both J-SBD and SBD were more than 95% pure (Fig. 2A, inset) and were folded, as indicated by their CD spectra (Fig. 2A). The CD spectrum of each protein indicated they were predominantly α-helical, with minima approximately at 208 and 222 nm. However, the J-SBD contained a relatively higher amount of α-helices than SBD alone (30% versus 10%), likely due to the presence of the J-domain which is predominantly αhelical, as shown by NMR studies with the *E. coli* J-domain (Pellecchia et al, 1996), in which approximately 49 of 76 residues are involved in formation of α-helices, while the SBD is formed by a mixture of α-helices and β-sheets (Fig. 1A).

**Fig. 2:**
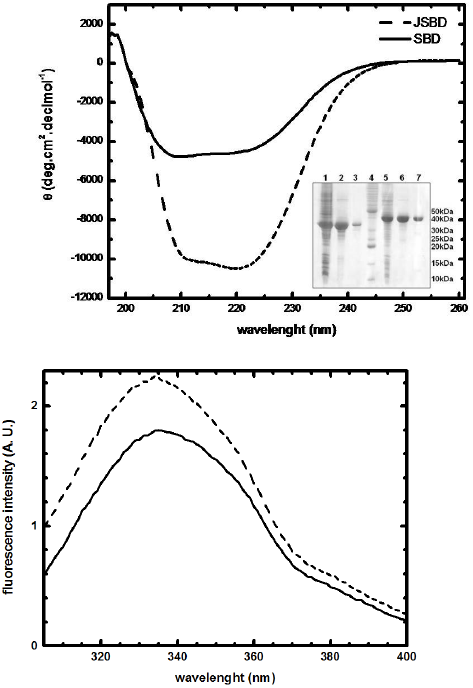
Protein purification and conformational measurements. **A)** Circular dichroism. Residual molar ellipticity [θ] was measured from 200 to 260 nm at 20 °C with protein solutions in 25 mM Tris-HCl (pH 8.0) buffer containing 150 mM NaCl. The spectra of J-SBD (dotted line) and SBD (solid line) are characteristic of α-helical rich proteins. Inset: SDS-PAGE, lane 1-3, soluble fraction of bacterial lyses, partial purification from affinity chromatography and elution from gel filtration chromatography of SBD, respectively; lane 4, molecular mass marker; lanes 5-7, soluble fraction of bacterial lyses, partial purification from affinity chromatography and elution from gel filtration chromatography of J-SBD, respectively. **B)** Fluorescence. Emission fluorescence spectra of J-SBD (dotted line) and SBD (solid line) at 20 °C measured with excitation at 295 nm and emission from 304 to 400 nm in 25 mM Tris-HCl (pH 8.0) buffer containing 150 mM NaCl. A.U., arbitrary units.

Emission fluorescence spectroscopy of Trp is a powerful technique to access information about the environment where this residue is located because of its sensitivity to the polarity of the environment. Therefore, emission fluorescence spectroscopy of Trp can be used to access the folded state of a protein (Royer, 2006). There is only one Trp in the SBD and none in the J-domain (Fig. 1B), thus this technique is useful to test whether or not the presence of the J-domain leads to conformational changes in the region of the Trp residue when experiments are performed with the same set up. The maximum fluorescence emission was 335 nm with a <λ> at 343 nm for both SBD and JSBD but the intensity of fluorescence of J-SBD was approximately 25% higher than that of the SBD alone at 335 nm (Fig. 2B and Table 1). These results suggested that, in both proteins, the Trp residue is buried in the hydrophobic core. However, although the Trp residue experienced the same environment polarity in each protein, there might be residues near the Trp in SBD that are partially quenching the fluorescence, which is released when the J-domain is present. Therefore, the addition of the J-domain causes subtle changes in the microstructure around the Trp residue.

**Table 1.** Summary of results.

| **EXPERIMENT** | **SBD** | **J-SBD** |
| --- | --- | --- |
| **CD (molar ellipticity [θ] at 222 nm)** | **-4400** | **-10500** |
| **FLUORESCENCE (λ<sub>max</sub>) nm** | **335** | **335** |
| **FLUORESCENCE (intensity relative to SBD at 335 nm)** | **1.00** | **1.25** |
| **TOM70 BINDING** | **Yes** | **Yes** |
| **OLIGOMERIC STATE** | **Multiple oligomeric species** | **Mainly monomeric** |
| **LUCIFERASE AGGREGATION PROTECTION** | **No** | **50% (at 1:2.5)** |
| **BIS-ANS BINDING DECREASED IN THE PRESENCE OF PEPTIDE** | **No** | **Yes** |
| **PROTEOLYSIS BY TRYPSIN** | **No** | **Yes** |

Another indication that J-SBD was folded and functional was that the protein showed the ability to bind the cytosolic fragment of Tom70, which is part of the translocator system in mitochondria. Tom70 is a TPR-domain protein that binds to Hsp70 thru its EEVD motif at the C-terminus (Young et al, 2003). The J-SBD chimera was able to bind Tom70, as shown by a pull-down assay in a similar way to the isolated SBD (Fig. S1).

### 3.2. The chimeric J-SBD was a monomer

Proteins from the Hsp70 family tend to oligomerize, a characteristic stimulated by temperature. Non-specific aggregates are formed at temperatures higher than 42 °C, and both high protein concentration and salt concentration (Carlino et al, 1992; Kim et al, 1992; Fouchaq et al 1999). The phenomenon of self-association has been reported for both DnaK, from *Escherichia coli,* and Hsc70s, from both bovine and human, and seems to be a conserved characteristic in evolution (Malinverni et al, 2015). The domain of Hsc70 that is responsible for oligomerization is the SBD, as the NBD alone does not oligomerize, while the SBD forms oligomeric species (Benaroudj et al 1997). Thus, we investigated whether the presence of the J-domain affects self-association. First, SECMALS was used to investigate the oligomeric state and molecular mass of the proteins (Fig. 3A). J-SBD eluted mainly as a single species with a molecular mass of 38.0 ± 2.0 kDa (Fig. 3A), which is in very good agreement to the molecular mass values of 40304.26 Da expected from its amino acid sequence (Fig. 1B). On the other hand, SBD alone eluted as multiple oligomeric species (Fig. 3A, inset).

**Fig. 3:**
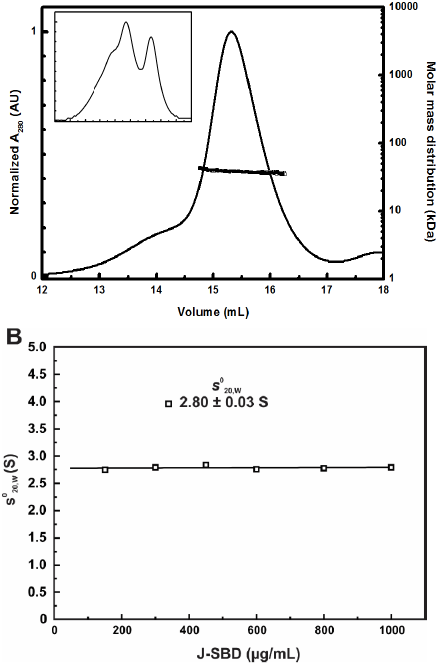
J-SBD is a monomer with molecular mass of approximately 38 kDa. **A)** SECMALS. The oligomeric state and molecular mass were investigated by size exclusion chromatography coupled with multi-angle laser light scattering (SEC-MALS). J-SBD eluted mainly as a single species with a molecular mass of 38.0 ± 2.0 kDa, a value in very good agreement with the molecular mass values of 40304.26 Da expected based on the amino acid sequence. Inset: SBD profile showing several species with molecular masses higher than that of the monomer. The experiments were performed using the Superdex 200 10/300 GL column (GE Life Sciences) coupled to a miniDAWN TREOS detector (Wyatt Technologies). **B)** Sedimentation velocity analytical ultracentrifugation data were fitted by the SedFit software to determine the coefficient sedimentation s_20,w_ from the c(S) distribution (see supplemental Fig. S1). *s*_20,w_ versus protein concentration was fit with a linear regression to calculate the *s*_0_^20,w^. This procedure minimizes errors caused by temperature, solution viscosity, and molecular crowding. Conditions: 35000 rpm, 20 °C, with J-SBD concentrations ranging from 150 to 1000 μg/mL in 25 mM Tris-HCl (pH 8.0) buffer containing 150 mM NaCl.

Next, sedimentation velocity AUC experiments were performed to investigate the oligomeric state and molecular mass of J-SBD. Data were analyzed by the SedFit software for c(S) distribution, showing a predominant species (>98% of the area under curve) with a peak centered at approximately 2.8 S at all protein concentrations tested (Fig. S2). The plot of s_20,w_ versus protein concentration (Fig. 3B) showed that J-SBD had a s^0^_20,w_ of 2.80 ± 0.03 S. From this result and fitting using SedFit software, J-SBD behaved as a monomer of 39 ± 1 kDa, again in very good agreement with the value measured from SEC-MALS and estimated by its amino acid sequence (see above). The fitting also suggested that J-SBD had an asymmetric shape, since the ƒ/ƒ_0_ calculated by the ratio of that maximum s-value by the s^0^_20,w_ was of approximately 1.5. Taken together, these results suggest that the J-SBD was an asymmetric monomer in solution.

### 3.3. The chimeric J-SBD had chaperone activity

The ability of J-SBD to protect a model protein from aggregation was investigated (Fig. 4). When heated at 42 °C, luciferase aggregated, as measured by an increase in signal at 320 nm (Fig. 4). Addition of increased concentrations of J-SBD, with stoichiometries from 1:1 to 1:10, partially protected luciferase from aggregation as measured by a decrease in signal at 320 nm (Fig. 4). At stoichiometry of 1:1.25 there was a decrease of approximately 50% in the measured signal. As controls, neither SBD nor a dummy protein, bovine serum albumin (BSA), had a significant effect on luciferase aggregation (data not shown), confirming that J-SBD has a specific effect in protecting luciferase. This result is striking because there are several pieces of experimental evidence suggesting that isolated Hsp70 does not protect luciferase against aggregation but does so only in the presence of Hsp40. For instance, Deloche et al. (1997) showed that DnaK alone, even at 2:1 stoichiometry, is not able to prevent Gdm-Cl unfolded luciferase from aggregation, whereas the addition of DnaJ, or its mitochondrial *Saccharomyces cerevisiae* orthologue, increased the amount of soluble luciferase from 10 to about 80%. Thus, the SBD appears to be interacting with the J-domain in the JSBD chimera and responding to it.

**Fig. 4:**
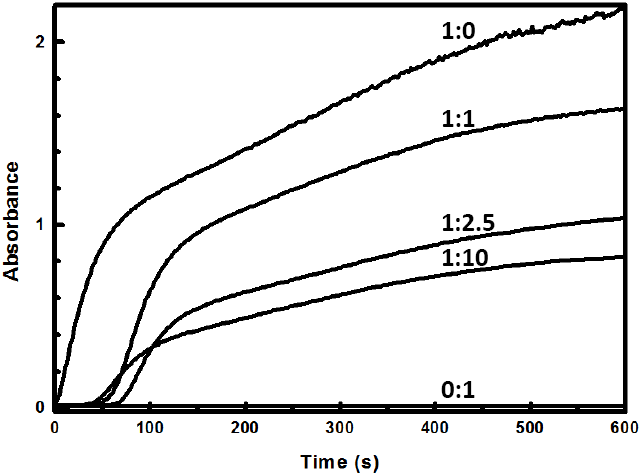
J-SBD protects luciferase from aggregation. Luciferase (1μM) alone (1:0), or in the presence of J-SBD (1:1-10), was heated at 42 °C and turbidity was followed by measuring at 320 nm. Buffer was 25 mM Tris-HCl (pH 8.0) buffer containing 150 mM NaCl. Luciferase aggregates in this condition (1:0), but it was protected by J-SBD, which did not aggregate (0:1).

Next, the capability to bind a model peptide was tested. Bis-ANS is a probe that is virtually non-fluorescent in aqueous solution or in the presence of a well-folded and compact protein, but it becomes fluorescent when bound to partially unfolded structures (Stryer, 1965). Figure 5 shows the intensity of emission fluorescence at 495 nm as a function of increasing concentrations of bis-ANS relative to both SBD and J-SBD at 25 °C. Both proteins bind bis-ANS, indicating that they contain exposed hydrophobic patches. J-SBD bound more bis-ANS than SBD on a molecule per molecule basis, which indicated that it had more exposed hydrophobic patches than SBD alone. The experiments were then performed in the presence of the mimetic peptide NRLLLTG, a model substrate for Hsp70, and the results clearly indicated that the binding of bis-ANS decreased when J-SBD was incubated with the peptide, whereas no effect was observed for SBD alone (Fig. 5). These results are in good agreement with a more compact conformation of SBD, which appears to resemble a ‘closed’ state. This is also corroborated by the fact this polypeptide was more resistant to trypsin digestion than JSBD (Fig. S3), even though the latter has ~50% more sites (47 versus 32).

**Fig. 5:**
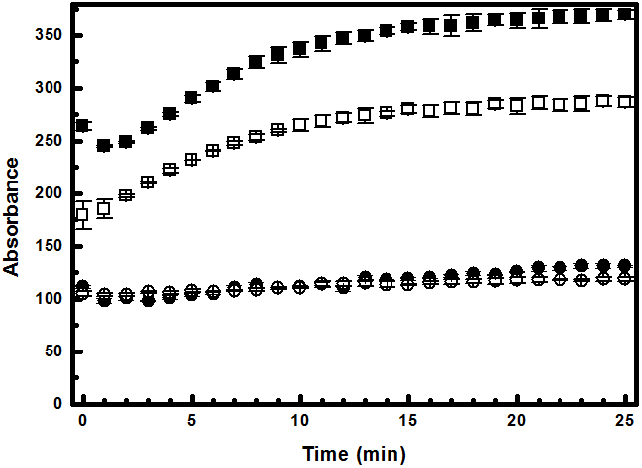
J-SBD binds the mimetic peptide (NRLLLTG). Intensity of emission fluorescence at 495 nm as a function of increasing concentrations of bis-ANS (tenfold) to SBD (circle) and J-SBD (square) at 25 °C. Experiments were performed in the absence (open) or in the presence (closed) of the peptide NRLLLTG, a model substrate for Hsp70. Binding of bis-ANS decreased when J-SBD was incubated with the peptide, whereas no effect was observed for SBD alone.

### 3.4. Final discussion and conclusions

The conformation and function of the chimeric J-SBD and isolated SBD were investigated, and the results are summarized in Table 1. Both proteins were produced folded with a high amount of secondary structure. Hydrophobic patches are partially exposed, as shown by the ability to bind bis-ANS, and the single Trp residue, located at the SBD, is buried in the apolar interior of the proteins. These results, together with the facts that the J-SBD has a larger amount of α-helical content due to the J-domain and that the SBD maintains its ability to bind a TPR-protein (TOM70), indicated that both the J-domain and the SBD are well-structured in the chimera and maintained their original conformation. Therefore, the chimeric construction, which is a fusion of the J-domain and the SBD in a single polypeptide, is a good model to promote the natural interaction of these domains, facilitating this investigation. However, the chimeric J-SBD showed properties not present in the SBD alone.

The self-association property of Hsp70 has been known for a long time and is a property of the SBD. Accordingly, the isolated SBD studied here showed this property but not the chimeric J-SBD. This chimera remained a monomer in all conditions tested, suggesting that the presence of the J-domain prevents the SBD from self-association. This result is in good agreement with a more recent model of the interaction of Hsp70 and Hsp40. In this view, the shift between oligomeric and monomeric states of Hsp70 is part of the cycle and is modulated by several factors, for instance, ATP, post-translational modification and Hsp40. One example is given in the work of Sarbeng et al. (2015), showing that mutants which affect Hsp70 dimerization decrease the affinity for Hsp40, and suggests a model in which Hsp40 binds to the dimeric Hsp70 presenting the ‘client’ protein and concomitantly induces both the monomerization and ATP hydrolysis of Hsp70. The chimera studied here has only specific domains of the entire Hsp70/Hsp40 complex, but the results clearly suggest that Hsp40 somehow influences the oligomeric state of Hsp70. The results suggest that, at some point in the cycle, interaction between the J-domain and the SBD favors the monomeric state of Hsp70.

Another interesting insight from the results with the chimeric J-SBD is that the presence of the J-domain induces the chaperone activity of the SBD. The chimeric JSBD showed fairly good protection against luciferase aggregation, while the SBD alone has only very minor activity. Additionally, there is good indication that the hydrophobic site in J-SBD that was available for bis-ANS was protected when in the presence of the peptide, indicating that it binds to J-SBD but not to SBD alone. In this model, the presence of the J-domain apparently induces an open conformation in the SBD, allowing for substrate binding.

In conclusion, this work collects suitable evidence for the action of the J-domain on the conformation and function of SBD, allowing us to suggest a model for the interaction of Hsp70 and Hsp40 (Fig. 6). In this model, the interaction stimulates monomeric Hsp70 with a ‘opened’ conformation that facilitates substrate binding. Of course, one has to take in consideration that the chimera has only one domain from each protein and thus cannot, of course, represent the interaction between the other domains. However, this model is likely a very good representation of one step in the complex cycle that is the interaction between Hsp70, Hsp40 and substrate proteins.

**Fig. 6:**
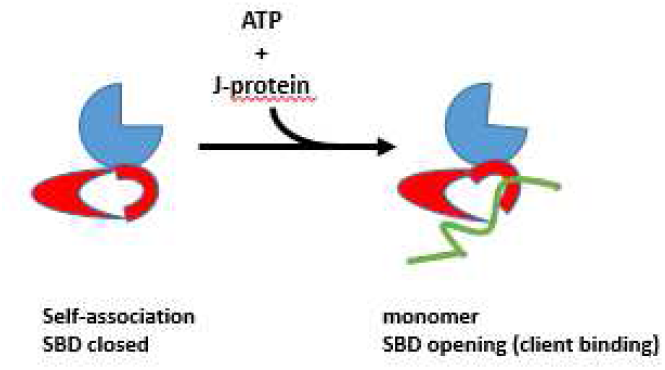
Model for the interaction between Hsp70 and Hsp40. In this model, the interaction stimulates a monomeric Hsp70 state with a ‘opened’ conformation that facilitates substrate binding.

## Acknowledgements

Research in the laboratory of CHIR is supported by grants from Fundação de Amparo a Pesquisa do Estado de São Paulo (FAPESP, 2012/50161-8), Conselho Nacional de Pesquisa e Desenvolvimento (CNPq) and CAPES. J.C. Borges thanks FAPESP for financial support (2014/07206-6 and 2017/07335-9) and CNPq for the Research Fellowship. We thank all LNBio technicians for their assistance.

